## Supplementary Data for "Occupancy estimation of wild species in a palm oil plantation using unstructured data"

Explanation of file data_for_repository.csv

This file contains the presence records collected in KAL, after cleaning and data augmentation. Only records from the classes Mammalia, Reptilia, Aves, Insecta and Magnoliopsida.

Meaning of columns.

StandardizedlocationID: name of the block

Sites code: code of the block

Species code: code of species

Latin name: name of species

23_species_in_analysis: 1 in case of a study species, else 0

Class: species group

Year: calendar year of observation

Habitat_of_site: main habitat of the block

Distance_to_large_forest: see text for explanation

Site size: area of block

List length: number of species (of all species groups, i.e., not only the five groups mentioned) Habitat_of_observation: habitat in which observation was made

Observer_name and observercode: speaks for itself

All_visits_made_in_one_year: Indicates whether all visits were made in only 1 year, and which year that is. This information was used to create alternative datasets (see text for explanation).

Observation: 1; there are only records in this file.

Explanation of file KAL_observers.csv

This file indicates for each observer how many records were collected from Mammalia, Reptilia, Aves, Insecta, and Magnoliopsida, respectively. This information was used to create alternative datasets (see text for explanation).
